# A systematic dissection of human primary osteoblasts *in vivo* at single-cell resolution

**DOI:** 10.1101/2020.05.12.091975

**Authors:** Yun Gong, Junxiao Yang, Xiaohua Li, Cui Zhou, Yu Chen, Zun Wang, Xiang Qiu, Ying Liu, Huixi Zhang, Jonathan Greenbaum, Liang Cheng, Yihe Hu, Jie Xie, Xuecheng Yang, Yusheng Li, Yuntong Bai, Yu-Ping Wang, Yiping Chen, Li-Jun Tan, Hui Shen, Hong-Mei Xiao, Hong-Wen Deng

## Abstract

Osteoblasts are multifunctional bone cells, which play essential roles in bone formation, angiogenesis regulation, as well as maintenance of hematopoiesis. Although both *in vivo* and *in vitro* studies on mice have identified several potential osteoblast subtypes based on their different transition stages or biological responses to external stimuli, the categorization of primary osteoblast subtypes *in vivo* in humans has not yet been achieved. Here, we used single-cell RNA sequencing (scRNA-seq) to perform a systematic cellular taxonomy dissection of freshly isolated human osteoblasts. Based on the gene expression patterns and cell lineage reconstruction, we identified three distinct cell clusters including preosteoblasts, mature osteoblasts, and an undetermined rare osteoblast subpopulation. This novel subtype was mainly characterized by the nuclear receptor subfamily 4 group A member 1 and 2 (NR4A1 and NR4A2), and its existence was confirmed by immunofluorescence staining. Trajectory inference analysis suggested that the undetermined cluster, together with the preosteoblasts, are involved in the regulation of osteoblastogenesis and also give rise to mature osteoblasts. Investigation of the biological processes and signaling pathways enriched in each subpopulation revealed that in addition to bone formation, preosteoblasts and undetermined osteoblasts may also regulate both angiogenesis and hemopoiesis. Finally, we demonstrated that there are systematic differences between the transcriptional profiles of human osteoblasts *in vivo* and mouse osteoblasts both *in vivo* and *in vitro*, highlighting the necessity for studying bone physiological processes in humans rather than solely relying on mouse models. Our findings provide novel insights into the cellular heterogeneity and potential biological functions of human primary osteoblasts at the single-cell level, which is an important and necessary step to further dissect the biological roles of osteoblasts in bone metabolism under various (patho-) physiological conditions.

## Introduction

Osteoblasts, which account for 4-6% of resident cells in bone, are derived from pluripotent bone marrow mesenchymal stem cells (BMMSCs) through the activation of signaling pathways regulated by osterix (OSX) and Runt-related transcription factor 2 (Runx-2)^1-4^. These cells are primarily known for their bone-building functions, including deposition of calcium phosphate crystals (e.g. hydroxyapatite), production of bone matrix constituents (e.g. type I collagen), as well as their ability to secrete a number of important proteins for bone metabolic processes such as integrin-binding sialoprotein (IBSP), secreted phosphoprotein 1 (SPP1), and bone gamma-carboxyglutamic acid-containing protein (BGLAP)^5,6^. In general, osteoblasts play a crucial role in the mineralization of the bone matrix^7,8^.

Many osteoblasts ultimately differentiate into osteocytes, which become embedded in the bone matrix to form a communication network for the regulation of bone formation and resorption. The osteoblasts and osteocytes regulate the differentiation of osteoclasts, which are primarily involved in bone resorption activities^9^. For instance, osteoblasts can promote osteoclast proliferation by producing macrophage-colony stimulating factor (M-CSF)^10,11^. Osteoblasts can also produce the receptor activator of nuclear factor kappa-B ligand (RANKL) and osteoprotegerin (OPG) to modulate osteoclast proliferation through the RANKL/RANK/OPG pathway^12^. On the other hand, osteoclasts may also regulate bone formation by osteoblasts^13^. The complex dynamics between the major bone cells control the delicate balance of bone formation/resorption that is critical for maintaining bone health.

While osteoblasts are typically associated with bone remodeling processes, previous studies have demonstrated that they also have the ability to interact with immune cells via the regulation of hematopoietic stem cell (HSC) niches. Specifically, primary osteoblast lineage cells synthesize granulocyte colony-stimulating factor (G-CSF), granulocyte M-CSF (GM-CSF), IL-1, IL-6, lymphotoxin, TGFß, TNFα, leukemia inhibitory factor (LIF), and stem cell factor (SCF); all of which play crucial roles in hematopoiesis^14,15,24,16–23^. Additionally, osteoblasts are responsible for modulating angiogenesis, which supports the high degree of vascularization in the bone marrow environment to provide sufficient oxygen for bone metabolism^25^.

Cellular heterogeneity is an essential characteristic of tissues of various cell populations. Although every cell shares almost the same genome, each cell acquires a unique identity and thus specific functional capabilities through molecular coding across the DNA, RNA, and protein conversions^26^. Therefore, even for the same known classical cell types, cells may be further classified into distinct subpopulations due to systematic differences in their gene expression profiles. Emerging evidence from *in vitro* studies in mice has revealed notable cell-to-cell heterogeneity within the osteoblast cell population. For instance, one study performed the soft agarose cloning technique on rat osteoblastic cells and detected diverse gene expression patterns in osteoblast cells at different stages of cellular differentiation^27^. Another study showed that the expression levels of osteoblastic specific markers including osteopontin, bone sialoprotein, and osteocalcin, varied in the mature osteoblasts of mice with different cellular morphology, suggesting that even these terminally differentiated osteoblasts were composed of multiple subgroups instead of a single unique cell group^28^. While these early efforts revealed the existence of osteoblast heterogeneity, the functional differences between distinct osteoblast subtypes were not well characterized.

The recent development of the state-of-the-art single-cell RNA sequencing (scRNA-seq) technology is expected to provide the most powerful approach to study the nature and characteristics of cell-to-cell heterogeneity. Compared with the conventional bulk RNA-seq approaches, scRNA-seq can reveal complex and rare cell populations, track the trajectories of distinct cell lineages in development, and identify novel regulatory relationships between genes^29^. Accumulating evidence from a few early scRNA-seq studies *in vivo* in mice has demonstrated the existence of several osteoblast subtypes. Baryawno et al.^30^ identified pre- and mature osteoblast subpopulations based on their transcriptional profiles, while Tikhonova et al.^31^ classified the osteoblasts into three subgroups and revealed significant changes in subgroup proportions under hematopoietic stress conditions induced by chemotherapy treatment. Although our understanding of osteoblast heterogeneity has evolved substantially based on these studies, the cell subtype characteristics of primary osteoblasts *in vivo* in humans has not been successfully explored. Studying the cellular heterogeneity in mice, or even the cultured cells from humans (though not yet existing), although useful, may not be ideal for studying human disease etiology. This is because the cell identity may vary between mice and humans, and cell culturing may systematically alter the gene expression of the studied cells^32^.

In this study, we successfully performed the first unbiased examination of the *in vivo* cellular landscape of freshly isolated human osteoblasts via scRNA-seq. We identified three distinct cell subtypes along with their differentiation relationships based on the transcriptional profiling of 5,329 osteoblast cells from the femur head of one human subject. We then compared the most differentially expressed genes (DEGs) of each cluster with known cell characterizing markers. Further, we identified distinct functional characteristics of each cell subpopulation suggesting their involvement in bone metabolism, angiogenesis modulation, as well as hematopoietic stem cell (HSC) niche regulation. The discovery of osteoblast subtypes is well beyond the scope of current gene expression studies for bone health and represents an important and necessary step to provide novel insights into bone physiological processes at the refined single-cell level.

## Results

### Osteoblasts identification

We applied an established protocol for *in vivo* human osteoblast isolation^33^ to obtain the alkaline phosphatase (ALPL)^high^/CD45/34/31^low^ cells from the femur head-derived bone tissue of one human subject (31-year-old Chinese male) with osteoarthritis and osteopenia through fluorescence-activated cell sorting (FACS). Several studies^34,35^ have demonstrated that this isolation protocol can successfully recover a high proportion of osteoblasts based on the elevated expression levels of osteoblastic markers via quantitative polymerase chain reaction (qPCR) or bulk RNA sequencing. In total, 9,801 single cells were encapsulated for cDNA synthesis and barcoded using the 10x Genomics Chromium system, followed by library construction and sequencing (**Figure 1A and Figure S1A**). After quality control, we obtained 8,557 cells, with an average of 2,365 and median of 2,260 genes detected per cell. A recently developed dimension reduction technique for scRNA-seq analysis, uniform manifold approximation and projection (UMAP)^36^, was applied to project the gene expression profiles on a two-dimensional panel for visualization of cellular heterogeneity (**Figure S1B**). After clustering the cells into six distinct subsets (C1-C6) by the k-nearest neighbor algorithm^37^, we used pairwise differential expression analysis for comparing each individual cluster against all the others to identify the DEGs of each subtype (**Figure S1C**).

**Figure 1.**
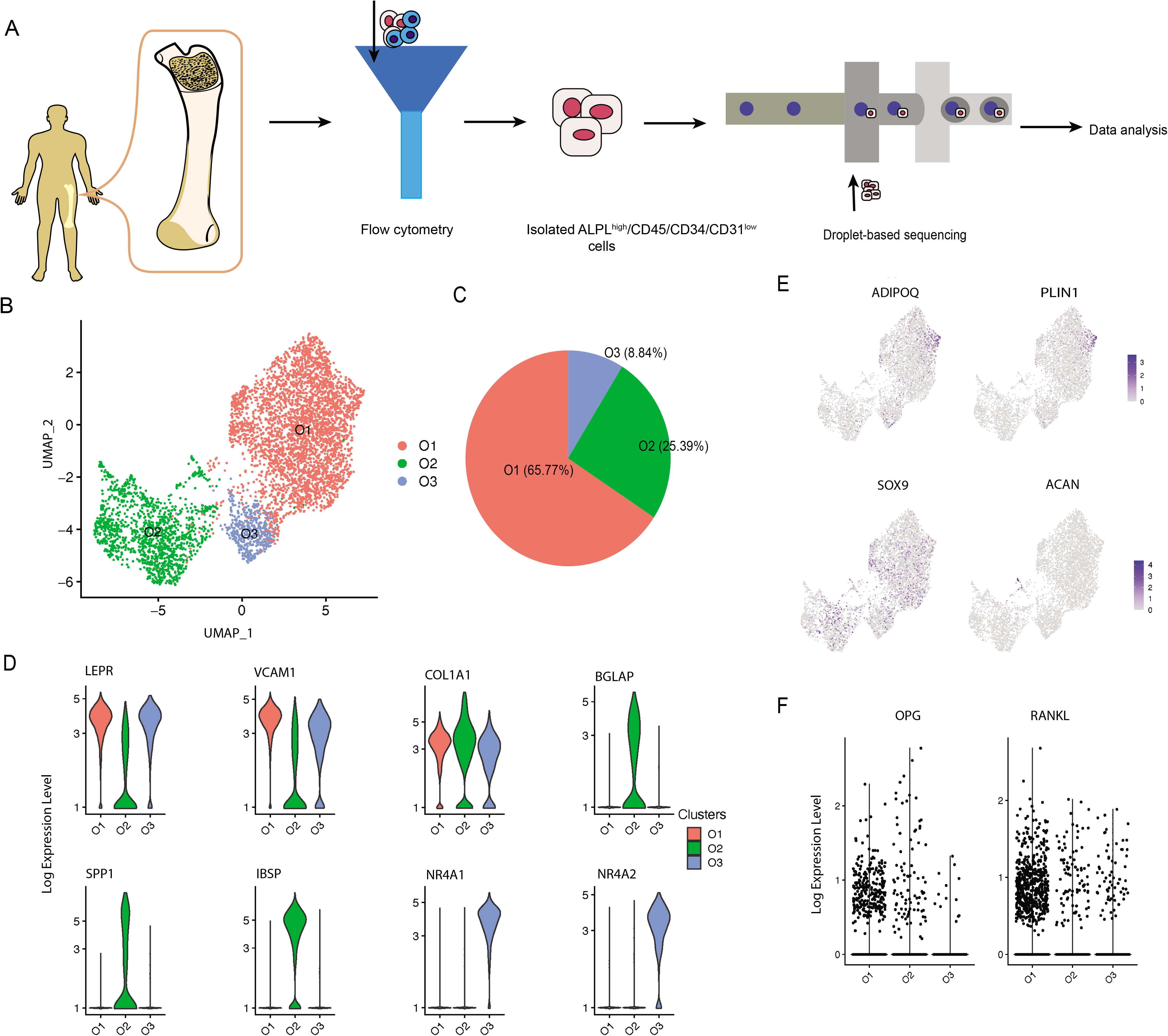
scRNA-seq Analysis of Human Osteoblasts. (A) Study overview. (B) Three osteoblast clusters. UMAP visualization of 5,329 osteoblasts, colored by clustering. (C) Proportion of three osteoblast clusters. Colored by clustering. (D) Cluster signature genes. Violin plots showing the log-transformed normalized expression levels of the two most significant marker genes in clusters O1, O2, and O3, respectively. (E) Log-normalized expression of adipocyte and chondrocyte biomarkers in osteoblast clusters. (F) Log-normalized expression of OPG and RANKL in three clusters.

While ALPL was enriched in clusters C1, C2 and C5, the osteoblast specific marker, RUNX2 (encoded by *RUNX2*)^32^, was only enriched in clusters C1 and C2, suggesting the presence of contamination by other cell types during the cell isolation process (**Figure S1D**). Based on the identified marker genes, the contaminant cells were classified as; 1) two nucleated erythrocyte groups (C3 and C4, expressing hemoglobin coding genes *HBB* and *HBA1*)^39^; 2) one smooth muscle cell group (C5, expressing smooth muscle alpha (alpha)-2 actin coding gene, *ACTA2*, and *CNN1*, which is specific to differentiated mature smooth muscle cells)^40,41^; and 3) one neutrophil group (C6, expressing neutrophil related genes *S100A8* and *MMP9*)^42,43^ (**Figure S1D**). These findings suggest that the protocol for human osteoblast isolation based on ALPL^high^/CD45/34/31^low^ may not be sufficient for specifically isolating osteoblasts alone, since ALPL is also highly enriched in other mesenchymal stem cell (MSC)-derived cells such as vascular smooth muscle cells^44^. Notably, by comparing the expression profiles between osteoblasts and contaminant cells, we found that the average fold change (avg_FC) of alpha-1 type I collagen (COL1A1, encoded by *COL1A1)* was large (avg_FC =4.497, **Table S1**), suggesting that positive selection based on the combination of ALPL and COL1A1 would be a more appropriate choice for human primary osteoblast isolation. While several studies^31,45^ in mice have used COL1A1 for osteoblast sorting, few (if any) studies have adopted this marker for human primary osteoblast isolation.

### scRNA-seq identifies multiple cell subtypes in human osteoblasts

To investigate the cellular diversity within osteoblasts, we extracted the clusters C1 and C2 with high expression levels of osteoblast related markers, i.e., RUNX2 and COL1A1 **(Figure S1D)**. After removing the cells with more than 5% of the transcripts attributed to mitochondrial genes, 5,329 high quality sequenced cells were retained for downstream analysis. Since the biomarkers of other BMMSC-derived cells such as adipocytes and chondrocytes were not enriched in the remaining cells^46^, we concluded that these cells are osteoblasts (**Figure 1E**). Based on the transcriptional profiles, we identified three cell subtypes of osteoblasts (**Figure 1B-1D**) which were annotated as; 1) O1 (65.77%), preosteoblasts, with relatively high expression levels of osteogenic BMMSC markers leptin receptor (encoded by *LEPR*)^47^ and vascular cell adhesion molecule 1 (encoded by *VCAM1*)^48^: 2) O2 (25.39%), mature osteoblasts, which showed highest expression levels of COL1A1 (*COL1A1*) and osteogenesis-associated genes including BGLAP (also known as osteocalcin, encoded by *BGLAP*), SPP1 (also known as osteopontin, encoded by *SPP1*), and IBSP (encoded by *IBSP)^49^*; 3) O3 (8.84%), undetermined osteoblasts, which not only expressed several BMMSC-associated genes (e.g., *LEPR* and *VCAM1*) but are also distinguished from the other subtypes by distinctively expressing high levels of nuclear receptor subfamily 4 group A member 1 and 2 (encoded by *NR4A1* and *NR4A2*). Notably, few osteoblasts in this cell population expressed OPG and RANKL (**Figure 1F**), which is consistent with the result proposed by Tat *et al*. that the expression levels of OPG and RANKL are significantly reduced in the osteoblasts from human osteoarthritic bone^50^

We further examined the transcriptional profiles of the three identified osteoblast subtypes and found that in addition to the cell markers, some bone development regulators were highly enriched in the pre-osteoblast and mature osteoblast clusters. For instance, preosteoblasts expressed insulin-like growth factor-binding protein 2 and 4 (IGFBP2 and IGFBP4, encoded by *IGFBP2* and *IGFBP4*) (**Figure 2A**). Previous studies have considered IGFBP2 as a stimulator for osteoblast differentiation through positive regulation of the AMP-activated protein kinase (AMPK)^51^, and demonstrated that IGFBP4 could stimulate adult skeletal growth in males^52^. Meanwhile, tenascin, a glycoprotein encoded by *TNC* that modulates osteoblast mineralization via matrix vesicles^53^, was highly enriched in mature osteoblasts. On the other hand, the cluster O3 showed high expression level of osteomodulin (OMD) (**Figure 2A**). It has been proposed that OMD induces endochondral ossification through PI3K signaling, which is an essential process for long bone formation^54^.

**Figure 2.**
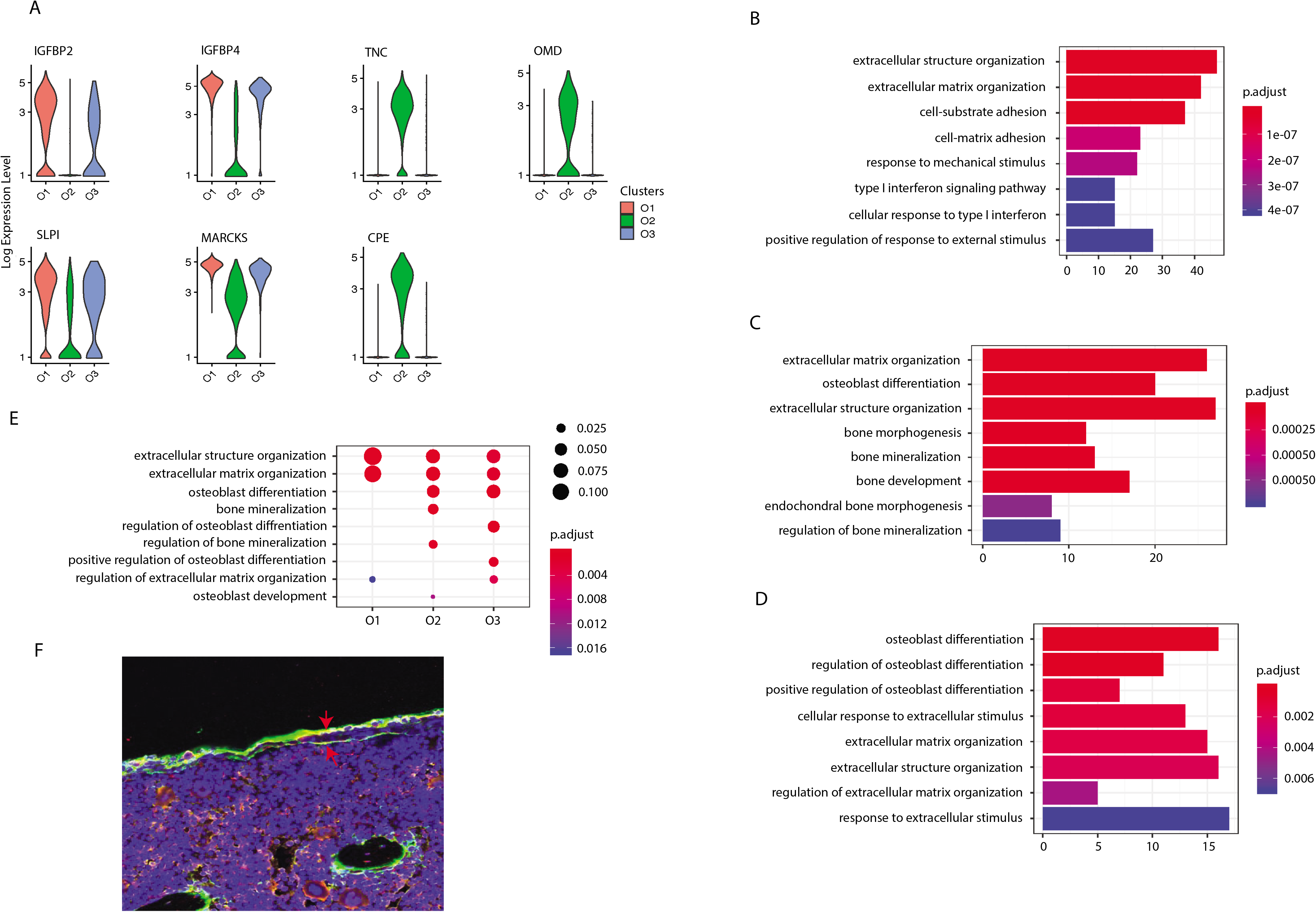
Osteoblast Subtypes and Cellular Functions in Bone Formation. (A) Osteoblasts related genes expressed in clusters O1, O2, and O3, respectively. (B-D) GO enrichment for the three osteoblast subpopulations, O1, O2, and O3, respectively. The length of the bar indicates the gene ratio (number of DEGs enriched in the GO term / total number of DEGs). The color indicates the adjusted p values for enrichment analysis. (E) Bone formation related GO terms enriched in clusters O1, O2, and O3. The size of dot indicates the gene ratio. The color indicates the adjusted p-value for enrichment analysis. (F) Immunofluorescence of mouse femur. The osteoblast marker ALPL was stained by green, while the cluster O3 marker NR4A1 was stained by red. The undetermined osteoblasts were located on the bone surface, co-stained by green and red (yellow).

In additional to the known osteoblast subtypes (pre-osteoblast and mature osteoblast), we also identified one potential novel osteoblast subtype (cluster O3), which is a major source for orphan nuclear receptors coding genes NR4A1 and NR4A2. Although previous studies have reported the presence of NR4A1 in mice osteoblasts induced by parathyroid hormone (PTH) *in vitro*^59^, the enrichment of this gene in osteoblasts *in vivo* has not yet been proposed or identified earlier in mice or in humans. It has been shown that NR4A1 and NR4A2 have an essential impact on the bone metabolism. Nuclear receptor subfamily 4 group A member 1 (NR4A1, encoded by *NR4A1*) has been proposed to inhibit osteoclastogenesis by producing an enzyme that breaks down the nuclear factor of activated T-cells (NFATc1) protein^60,61^. Additionally, the reduced expression of nuclear receptor-related 1 (NURR1, encoded by *NR4A2*) inhibits the expression of osteoblast differentiation marker genes such as *BGLAP* and *COL1A1^62^*. The immunofluorescence on mouse femur illustrated the co-staining of the osteoblast marker ALPL with the biomarker NR4A1 on the bone surface, thereby confirming the existence of the novel O3 cluster *in vivo* in humans (**Figure 2F**).

While all the osteoblast clusters highly expressed the osteoblastogenic gene *RUNX2*^38^, they may affect bone development from different aspects. To further investigate the distinct biological functions related to osteogenesis among the three clusters of osteoblasts, we performed gene ontology (GO) enrichment analysis based on the DEGs (avg_FC > 1.200, **Table S2**) in each cluster. All the clusters were enriched in GO terms related to bone development including “extracellular matrix organization” and “extracellular structure organization” (**Figures 2B-2E**). The preosteoblasts and mature osteoblasts are involved in the extracellular matrix (ECM) formation through production of different types of collagen, such as types 3, 14, and 18 in preosteoblasts and types 1, 12, and 13 in mature osteoblasts (**Tables S3** and **S4**). The mature osteoblasts were uniquely enriched for “bone mineralization”, while mature and undetermined osteoblasts were both enriched for “osteoblast differentiation” (**Figures 2C-2E, Tables S4** and **S5**). Additionally, the undetermined osteoblasts were uniquely enriched for “regulation of osteoblast differentiation” and “positive regulation of osteoblast differentiation” (**Figure 2D**). Surprisingly, few GO terms related to osteoblast differentiation are enriched in the preosteoblasts, suggesting that the undetermined and mature osteoblasts may modulate the osteoblast differentiation processes to a larger extent than preosteoblasts.

### Dynamic gene expression patterns reveal the differentiation relationship between different osteoblast subtypes

In order to reveal the differentiation dynamics of the osteoblast cell population, we reconstructed the developmental trajectory of the three identified clusters of osteoblasts. All the cells were contained within one cellular lineage without any bifurcations (**Figure 3A**), suggesting that only one terminal subtype exists in the osteoblastic population. While preosteoblasts and undetermined osteoblasts were highly enriched in the early stages of pseudotime, mature osteoblasts were distributed in the terminal stages of the osteoblastic lineage (**Figure 3A** and **C**). To strengthen the trajectory inference, we also assessed the transcriptional continuum of the cell lineage. The results showed that the expression of BMMSC associated genes, *LEPR* and *APOE*^63^, decreased over pseudotime, while the expression levels of osteogenic markers such as *RUNX2, ALPL, BGLAP, SPARC*^64^, and *COL1A1* were the highest at the end stages of pseudotime (**Figure 3B**). This result is consistent with the findings from other studies that have examined the transcriptional continuum in osteoblastic differentiation^65^. The analysis of the gene expression profiles of mice osteoblasts cultured *in vitro* showed that the expression levels of NR4A1 rapidly increased from day 5 to day 7 whereas the expression of ALPL remained relatively stable throughout the osteoblast development (**Figure 3D**), further supporting the conclusion that the undetermined osteoblasts are involved in the early stages of the osteoblastic lineage.

**Figure 3.**
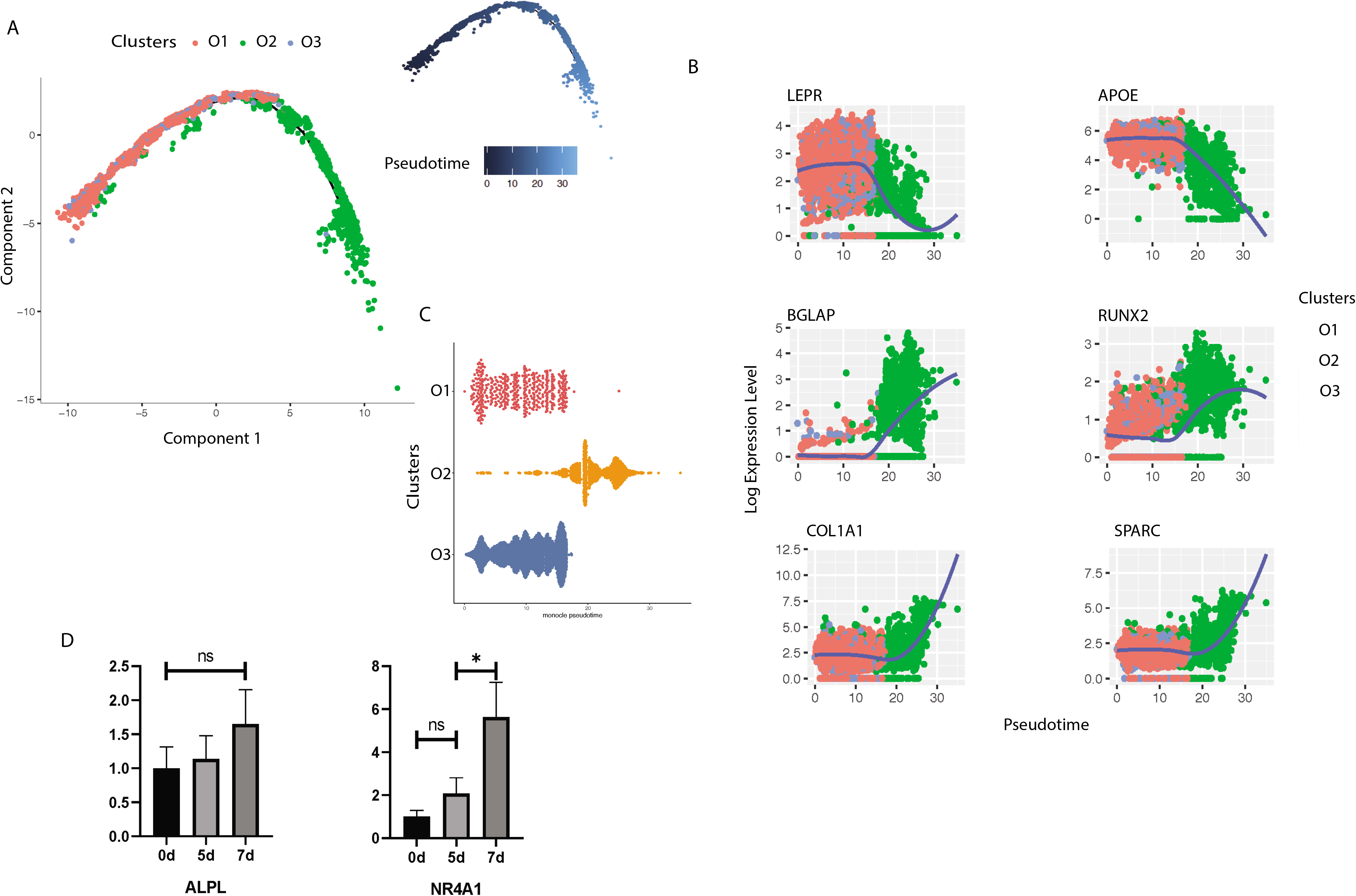
Trajectory Inference of Human Osteoblasts. (A) Reconstructed cell differentiation trajectory of human osteoblasts. The upper-right trajectory plot in the square indicates the direction of pseudotime. (B) Expression levels (log-normalized) of indicated genes in the three osteoblast subtypes with respect to their pseudotime coordinates. The x-axis indicates the pseudotime, while the y-axis represents the log-normalized gene expression levels. The color corresponding to the three different osteoblast subsets. The loess regression was applied to fit the relationship between pseudotime and expression level. (C) Cell distribution based on the pseudotime coordinates. The x-axis is the pseudotime and the y-axis represents the osteoblast subtypes. (D) Expression levels of ALPL and NR4A1 in mice osteoblasts *in vitro* at day 0, 5 and 7, respectively. N.S., not significant, *p-adjusted ≤ 0.05, **p-adjusted ≤ 0.01, *** p-adjusted ≤ 0.005.

### Preosteoblasts regulate angiogenesis through multiple signaling pathways

Although evidence has shown that osteoblasts can regulate angiogenesis^25,66^, few studies have explored this relationship at the single-cell level. Based on functional enrichment of the highly expressed genes, we found that the GO term “regulation of angiogenesis” was enriched in preosteoblasts but not in the other clusters (**Figure 4A**). We further investigated the enriched genes in preosteoblasts and found that 22 genes are involved in angiogenesis (**Table S3**). In particular, the preosteoblasts showed significantly higher expression levels of *SFRP1, MDK*, and *THBS1* compared with the other clusters (**Figure 4B**). It has been demonstrated that Secreted Frizzled-related protein-1 (sFRP1, encoded by *SFRP1*), a modulator of Wnt/Fz pathway, can modify MSC capacities enabling MSCs to increase vessel maturation^67^. Midkine (MDK, encoded by *MDK*) is an enhancer of angiogenesis, and MDK expression has been shown to be positively correlated with vascular density in bladder tumors^68^. Thrombospondin 1 (THBS1, encoded by *THBS1*) is considered to be a potent endogenous inhibitor of angiogenesis through antagonization of vascular endothelial growth factor (VEGF) activity^69^.

**Figure 4.**
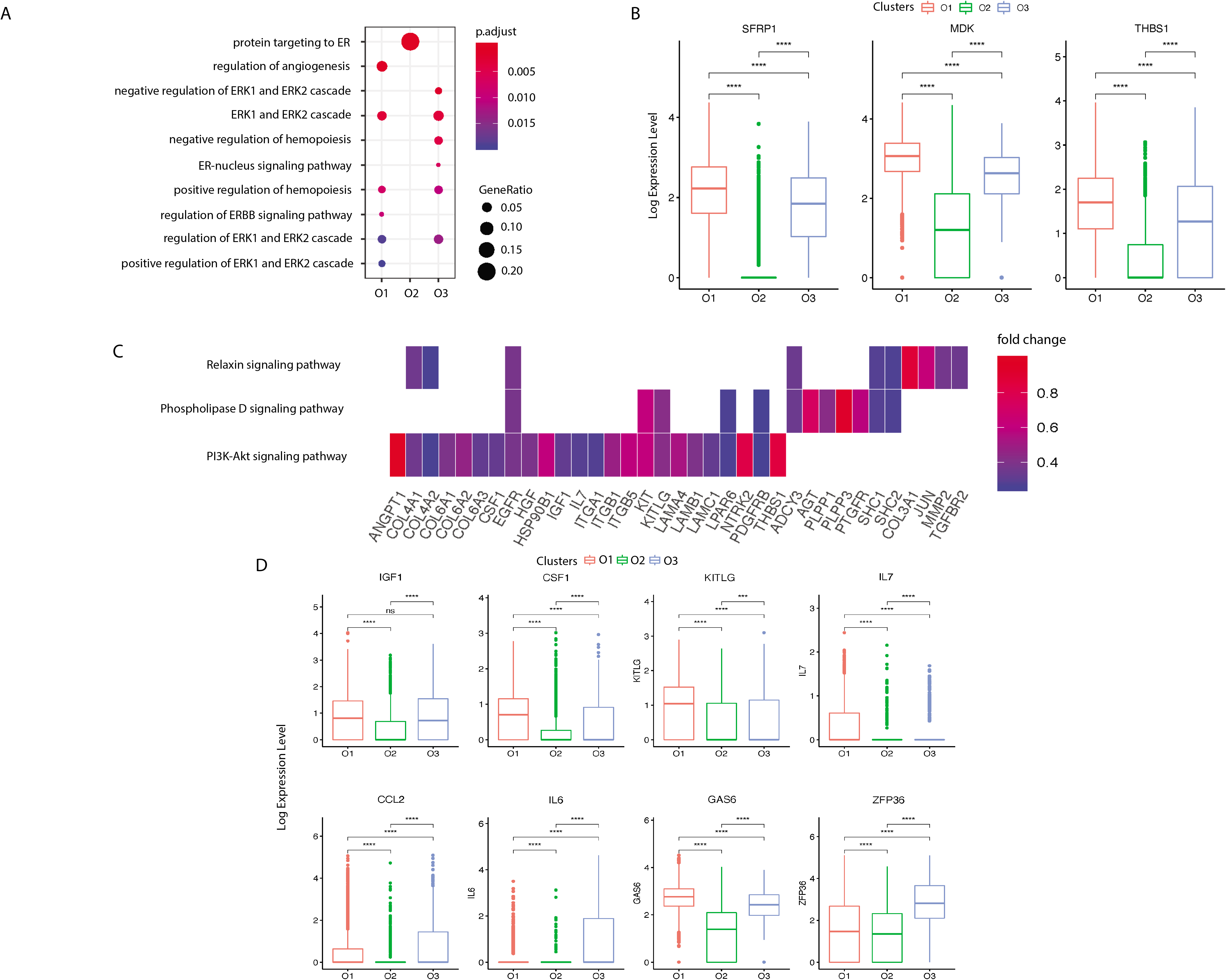
Pre- and Undetermined Osteoblasts Regulate Angiogenesis and Hemopoiesis. (A) Angiogenesis and hematopoiesis modulation related GO terms enriched in the three clusters. The x-axis represents the clusters and the y-axis is the GO terms related to the angiogenesis and hematopoiesis regulation. The size of the dot indicates the gene ratio and the color indicates the adjusted p-values. (B) Angiogenesis associated genes enriched in cluster O1. X-axis represents the three clusters and y-axis reflects log-normalized gene expression levels. The data are mean ± standard deviation. Stars indicate the significance levels of the gene expression difference between two clusters by Wilcoxon signed-rank test. N.S., not significant, *p-adjusted ≤ 0.05, **p-adjusted ≤ 0.01, *** p-adjusted ≤ 0.005. (C) Gene expression pattern in enriched pathways. Squares show enriched DEGs in the corresponding terms (rows). Color indicates the expression value of the DEGs (average logFC). (D) Hemopoiesis factors enriched in clusters O1 and O3.

Next, we investigated the signaling pathways enriched in preosteoblasts that are related to angiogenesis. Kyoto encyclopedia of genes and genomes (KEGG) pathway analysis revealed three highly enriched pathways including PI3K-Akt, phospholipase D, and relaxin signaling pathways, which are known to be implicated in angiogenesis modulation^70–72^ (**Figure 4C**). Several genes related to the “regulation of angiogenesis” GO term were enriched in these pathways (**Table S3**, **Figure 4C**), including *COL4A2, TGFBR2*, and *IL7*. These findings suggest that preosteoblasts may regulate angiogenesis to a larger extent than other osteoblast subtypes.

### Osteoblast populations modulate the development of HSC niche

Osteoblasts are parts of the stromal cell support system that provides critical regulators of hematopoiesis^73^. It has been reported that osteoblasts lining the bone surface can generate an extracellular environment supporting non-skeletal hematopoietic cells^74^. To identify the specific roles of each osteoblast subpopulation in hematopoiesis regulation, we first examined the expression patterns of hematopoiesis related factors in all the three osteoblast subtypes. We found that while both preosteoblasts and undetermined osteoblasts highly expressed insulin-like growth factor 1 (IGF-1, encoded by *IGF1*), which is a growth factor that has been shown to be responsible for burst-like growth of early erythroid progenitor cells *in vitro*^75^, the preosteoblasts also expressed high levels of M-CSF (encoded by *CSF1*), stem cell factor (SCF, encoded by *KITLG*), and interleukin 7 (IL-7, encoded by *IL7*) (**Figure 4D**). This suggests that preosteoblasts may play an important role in modulating macrophage development^76^, preventing HSC apoptosis^77^, and regulating T-cell differentiation^78^. We also found that the undetermined osteoblasts represent a major source of C-C motif chemokine ligand 2 (CCL-2, encoded by *CCL2*) (**Figure 4D**), which is a critical factor for monocyte recruitment in acute inflammatory response^79^. Additionally, the undetermined osteoblasts expressed interleukin 6 (IL-6, encoded by *IL6*), which displays a strong synergy with IL-7 to stimulate naïve CD8^+^ T cells^80^. Therefore, the undetermined osteoblasts may coordinate with preosteoblasts to support the hemopoietic niche. However, few hematopoietic factors were enriched in mature osteoblasts, suggesting that they may have limited impact on the hematopoietic system.

We investigated the predicted functions associated with hematopoiesis regulation enriched in the preosteoblasts and undetermined osteoblasts. Notably, while the GO term “positive regulation of hemopoiesis” was enriched in preosteoblasts, the highly expressed genes of the undetermined cluster were related to the negative regulation of hemopoiesis (**Figure 4A**). Apart from the growth factors mentioned above, preosteoblasts also had a significantly higher expression level of growth arrest-specific 6 (GAS6, encoded by *GAS6*) (**Figure 4D**), which can induce natural killer cell development via its positive regulatory effect on fibromyalgia syndrome-like tyrosine kinase 3 (FLT3) signaling in CD34+ hematopoietic progenitor cells (HPCs)^81^. Meanwhile, the undetermined subgroup had significantly higher expression levels of zinc-finger protein 36 (ZFP36, encoded by *ZFP36*) (**Figure 4D**), which can inhibit the erythroid differentiation^82^. These findings suggest that these two osteoblast subgroups may play different roles in hemopoiesis regulation, and that the relative proportions and functional levels of these subtypes within the osteoblast cell population may be crucial for maintaining the homeostasis of hematopoiesis.

### Transcriptional divergence of human and mouse osteoblasts

Due to the limited *in vivo* transcriptomics studies in mouse osteoblasts at different developmental stages, we first compared the expression profiles of osteoblasts acquired from humans *in vivo* and the osteoblasts from mice cultured *in vitro*. In addition, the *in vitro* mouse osteoblast differention data can clearly demonstrate the gene expression trajectory as compared to the inferred gene expression trajectory in *in vivo* mice data obtained from single cell sequencing. Several fundamental differences were observed. For instance, all the three human osteoblast clusters *in vivo* expressed secretory leukocyte protease inhibitor (encoded by *SLPI*) (**Figure 2A**), a serine protease inhibitor that promotes cell migration and proliferation while also suppressing the inflammatory response^55^. This conflicts with the previous *in vitro* findings in mice^56^, which demonstrated the enrichment of SLPI in mature but not in preosteoblasts. Additionally, we found that the human preosteoblasts isolated *in vivo* showed a significantly higher expression level of SLPI compared with mature osteoblasts (**Figure 2A**). In further contrast, the mature mouse osteoblasts culutured *in vitro* were highly enriched for myristoylated alanine rich protein kinase C (PKC) substrate (MARCKS, encoded by *MARCKS*), while this gene was highly expressed by the preosteoblasts but not the mature osteoblasts in humans *in vivo* (**Figure 2A**). It has been shown that *MARCKS* is the PKC-*δ* effector which modulates cathepsin K secretion and bone resorption in osteoclasts^57^. Remarkably, we also found that *CPE*, encoding carboxypeptidase E, was only enriched in mature osteoblasts *in vivo* in humans (**Figure 2A)**, which is inconsistent with the previous mouse study *in vitro* finding that the expression level of CPE decreases throughout osteoblast development^56^. While *CPE* plays an important role in RANKL-induced osteoclast differentiation^58^, it is plausible that a reduction in the proportion of mature osteoblasts could attenuate osteoclast proliferation.

To better understand the systematic differences in osteoblastic transcriptional profiles between human and mouse, we then integrated our sequencing data with the publicly available data from a previous scRNA-seq study^31^ on mouse osteoblasts *in vivo* (**Figure 5A**). After correcting for the potential batch effects between independent experiments using canonical correlation analysis (CCA)^37^, a moderate positive correlation (R = 0.53, p-value < 2.23e-16) was observed between the human and mouse osteoblastic transcriptomic profiles (**Figure 5B**). This suggests that while some common characteristics may exist between human and mouse osteoblasts, there are considerable differences in the gene expression profiles at the single-cell level.

**Figure 5.**
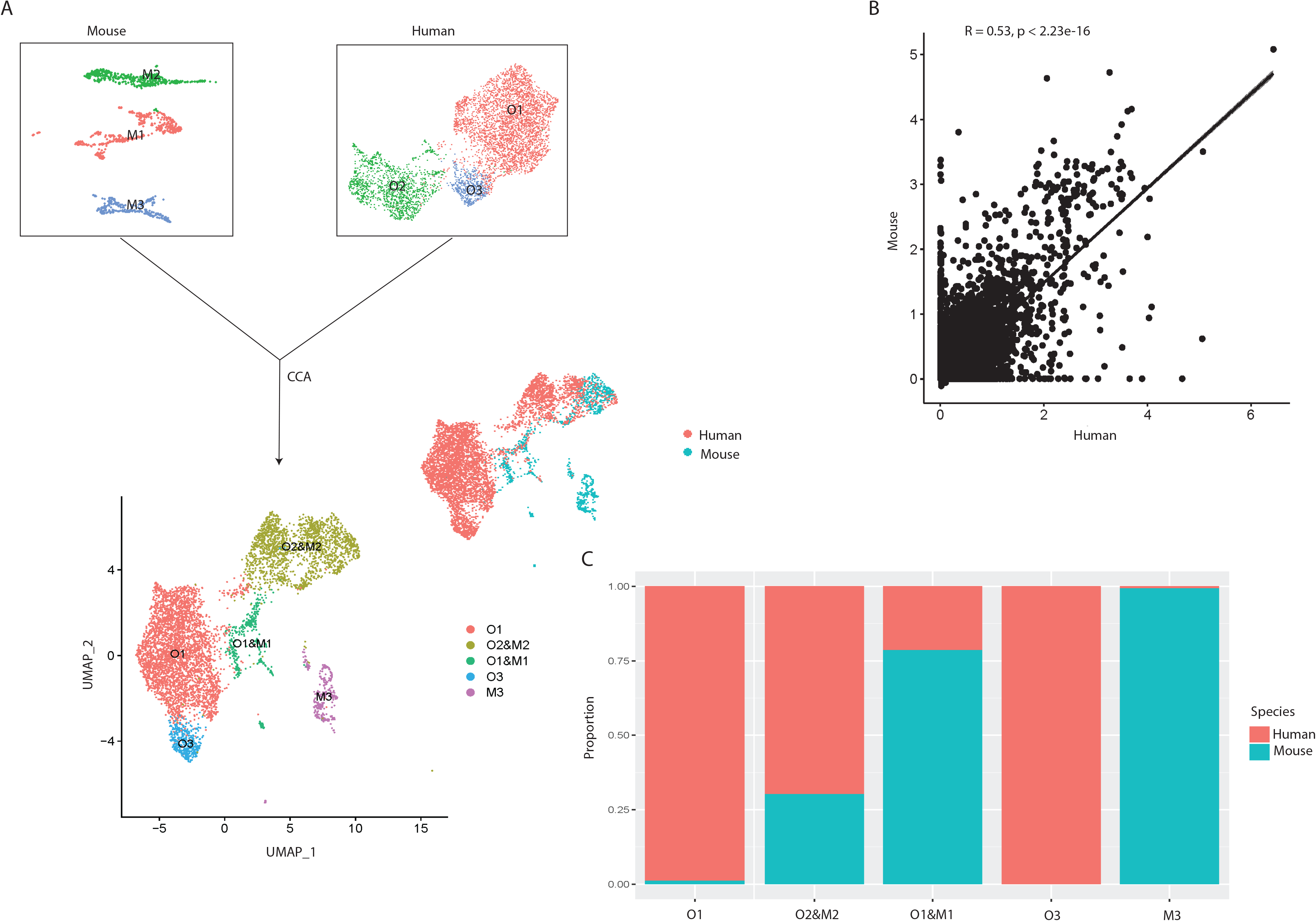
Integrated Cross-species Analysis Between Human and Mouse Osteoblasts. (A) UMAP visualization of human and mouse osteoblast integration. The upper-left plot represents the three osteoblast clusters in mice, the upper-right plot indicates the three osteoblast subtypes in humans. The bottom plot represents the five clusters after the integration, colored by clusters. The middle-right is the integrated data colored by different species. (B) Correlation of gene expression among different osteoblast datasets after CCA integration. Each dot represents an individual gene. Axes measure the average gene expression levels (logFC) in the indicated subject. Correlations were measured by Spearman correlation coefficients (*R*) (*p* < 0.01). (C) Proportion of the human and mouse osteoblasts in each cluster after CCA integration, colored by different species. The x-axis represents five different clusters and y-axis indicates the proportion.

To further investigate the shared and distinct features among human and mouse osteoblasts, we applied unbiased clustering analysis with UMAP^36^ for dimension reduction. The result showed that human osteoblast cluster O2 overlapped with mouse osteoblast subtype M2, which represents the mature osteoblasts in mice^31^ (**Figure 5A** and **5C**), indicating that the overall gene expression patterns of human and mouse mature osteoblasts are highly similar. Furthermore, we also found strong correlations in the expression patterns between small subgroups in human preosteoblasts (cluster O1) and the mouse osteoblast subtype M1^31^ (**Figure 5A** and **C**). In contrast, the human osteoblast cluster O3 did not overlap with any osteoblast subtypes in mice (**Figure 5A**). These findings suggest that the transcriptional profiles of human and mouse osteoblasts may demonstrate systematic differences in early cellular developmental stages but share similar features at the terminal stage of osteoblastic development lineage. Therefore, studies based on osteoblasts acquired from mouse models may introduce notable bias when inferring the biological characteristics of human osteoblasts, especially for those that have not yet reached a mature state.

## Discussion

While it is now well-appreciated that human osteoblast heterogeneity may be distinguished by differentiational stages^27^, the full spectrum of cells that comprise the osteoblast population, especially *in vivo* in humans, has remained elusive. In this study, for the first time, we classified the freshly isolated primary osteoblasts from human bone (without any *in vitro* culturing) into three subpopulations based on systematic differences in gene expression profiles and revealed their distinct functional roles in bone metabolism as well as in the regulation of angiogenesis and hemopoiesis. Further, in contrast to the results proposed by Tarkkonen *et al.^83^*, our results demonstrate systematic differences in the transcription profiles between humans and mice, emphasizing the importance of *in vivo* studies in humans.

Here, we highlight some of the key findings. First, our results indicate that the osteoblast isolation technique typically used in the field^33^, FACS isolation based on ALPL, was not sufficient for specific isolation of osteoblasts since the isolated cell population included approximately 40% contamination by the other cell types. By comparing the transcriptomic profiles between osteoblasts and contamination cells, we hypothesized that the combination of ALPL and COL1A1 would reduce contamination in osteoblast selection. In addition to known osteoblast subtypes, premature and mature osteoblasts, we also identified one rare osteoblast subpopulation which highly expressed NR4A1 and NR4A2. Based on the gene expression patterns and the inferred osteoblastic lineage trajectory, we found that: 1) Although both preosteoblasts (LEPR^high^/VCAM1^high^) and undetermined osteoblasts (NR4A1^high^/NR4A2^high^) are in the differentiation lineage ordering, preosteblasts are primarily responsible for ECM organization during bone formation processes as well as inducing hematopoiesis and modulating angiogenesis, while undetermined osteoblasts are mainly involved in the regulation of osteoblastogenesis and inhibition of hematopoiesis; 2) mature osteoblasts (IBSP^high^/BGALP^high^) arise at the terminal stages of cellular differentiation and are crucial for bone mineralization.

Despite the novelty of this scRNA-seq study in freshly isolated human osteoblasts, an important limitation is that all the cells were collected from the femur head of one 31-year-old Chinese male subject with osteoarthritis and osteopenia. This might introduce some bias in the osteoblast subpopulation identification and cell subtype proportion estimation compared with healthy individuals. Furthermore, due to limited cell numbers, the differentiation relationships between preosteoblasts and undetermined osteoblasts have not been clarified here. Future studies based on a larger sample size are needed to uncover the disease associated changes in cell subtype compositions, as this will have significant implications for the development of novel therapeutics. Despite these potential limitations, our results provide the first necessary and valuable insights into the cellular heterogeneity of osteoblasts along with a comprehensive and systematic understanding of the cell-intrinsic and cell-specific mechanisms which may underlie bone metabolism and associated disorders.

## Acknowledgement

This research was benefited by grants from the National Institutes of Health (R01AR069055, U19AG055373, P20GM109036, R01AG061917), National Natural Science Foundation of China (Grant No. 81902277), National Key R&D Program of China (Grant No. 2017YFC1001100), Natural Science Foundation of Hunan Province (S2019JJQNJJ2093), Changsha Science and Technology project (kq1907153), Central South University (Grant Nos. 164990007, 2018zzts886), and Xiangya Clinical Big Data Project (xyyydsj9).

## Author contributions

YG wrote the main manuscript text and conducted major analysis; JY and LC collected the human sample and corresponding clinical information; XL, YC, and CZ performed the experiments; JG, HS, YPW, YPC and HWD did language proofreading; ZW, LJT, and YB prepared supplementary information and validated the results; the study was conceived, designed, initiated, directed, and supervised by HS, HMX, and HWD. All authors participated in the discussions of the project and reviewed and/or revised the manuscript.

## Data availability

The scRNA-seq data for primary osteoblasts from one human sample can be accessed with accession number under GSE147390 in GEO database. One previous scRNA-seq data of mice osteoblasts used in this study can be accessed with accession number of GSE108891.

## Methods

### Study population

The clinical study was approved by the Medical Ethics Committee of Central South University and written informed consent was obtained from the study participant. The study population included one 31-year-old male with osteoarthritis and osteopenia (BMD T-score: 0.6 at lumbar vertebrae, −1.1 at total hip), who underwent hip replacement at the Xiangya Hospital of Central South University. The subject was screened with a detailed questionnaire, medical history, physical examination, and measured for bone mineral density (BMD) before surgery. Subjects were excluded from the study if they had preexisting chronic conditions which may influence bone metabolism including diabetes mellitus, renal failure, liver failure, hematologic diseases, disorders of the thyroid/parathyroid, malabsorption syndrome, malignant tumors, and previous pathological fractures^84^. The femur head was collected from the patient during hip replacement surgery. The specimen was shortly stored in 4°C and transferred to the laboratory within 2 hours, where it was then processed within 24 hours after delivery.

### Mice

Female C57BL/6J mice were purchased from Jackson Laboratory (Bar Harbor, ME, USA). All mice were housed in pathogen-free conditions and fed with autoclaved food. All experimental procedures were approved by the Ethics Committee of Xiangya Hospital of Central South University.

### BMD measurement

BMD (g/cm^2^) was measured using DXA fan-beam bone densitometer (Hologic QDR 4500A, Hologic, Inc., Bedford, MA, USA) at the lumbar spine (L1 −L4) and the total hip (femoral neck and trochanter) as described previously by our group ^85,86^. According to the World Health Organization definition^87^ and the BMD reference established for Chinese population^88^, subjects with a BMD of 2.5 SDs lower than the peak mean of the same gender (T-score ≤ −2.5) were determined to be osteoporotic, while subjects with −2.5 < T-score < −1 are classified as having osteopenia and subjects with T-score > −1.0 are considered healthy.

### Isolation of osteoblasts

Osteoblasts were extracted from the human femur head based on the widely used dissociation protocols^33^ with a few minor adjustments. Briefly, bone tissue samples were chopped into small fragments and washed twice with phosphate-buffered saline (PBS). These fragments, containing a mixture of cortical and trabecular bone, were then incubated with a highly purified, endotoxin-free type II collagenase (1 mg/ml, Cat: A004174-0001, Sangon Biotech, Shanghai, China) at 37.0°C for 30 mins and 60 mins in the first and second round of digestion respectively. After the incubation, the solution was filtered through a 70 μm filter and incubated with red blood cell lysis buffer (Cat: R1010, Solarbio Science & Technology CO., Beijing, China) for 5 mins. The collected cells were washed twice with PBS.

### Fluorescence-activated cell sorting (FACS) enrichment for osteoblasts

Before FACS, collected cells were incubated with human CD31/34/45-PE (Cat:303106, Cat:343606, Cat:304008, BioLegend, San Diego, CA, USA), human ALPL-APC (Cat: FAB1448A, R&D Systems, Minneapolis, MN, USA), and 7-AAD (488 nm) antibody (R&D Systems, Minneapolis, MN, USA). FACS was performed on a BD FACS Aria II sorter (Becton-Dickinson, Franklin Lakes, NJ, USA), where dead cells and debris were excluded by forward versus side scatter (FSC/SSC), and live cells were further enriched by negative selection of 7-AAD. ALP+/CD45/34/31-cells were collected as osteoblasts (**Figure S1**)^33^. Gating schemes were established with fluorescence-minus-one (FMO) controls.

### Single-cell RNA-seq (scRNA-seq) library preparation and sequencing

scRNA-seq libraries were prepared using Single Cell 3’ Library Gel Bead Kit V3 following the manufacturer’s guidelines (https://support.10xgenomics.com/single-cell-gene-expression/library-prep/doc/user-guide-chromium-single-cell-3-reagent-kits-user-guide-v3-chemistry). Single cell 3’ Libraries contain the P5 and P7 primers used in Illumina bridge amplification PCR. The 10x Barcode and Read 1 (primer site for sequencing read 1) were added to the molecules during the GEM-RT incubation. The P5 primer, Read 2 (primer site for sequencing read 2), Sample Index and P7 primer were added during library construction. The protocol is designed to support library construction from a wide range of cDNA amplification yields spanning from 2 ng to 2 μg without modification. All constructed single-cell RNA-seq libraries were sequenced on the Illumina Novaseq6000 platform with a sequencing depth of at least 100,000 reads per cell for a 150bp paired end (PE150) run.

### Pre-processing of single-cell RNA-seq data

We demultiplexed the cellular barcodes and aligned reads to the human transcriptome (GRCh38/hg38) using Cell Ranger 3.0 (https://support.10xgenomics.com/single-cell-gene-expression/software/pipelines/latest/what-is-cell-ranger). To create Cell Ranger-compatible reference genomes, the references were rebuilt according to instructions from 10x Genomics (https://www.10xgenomics.com), which performs alignment, filtering, barcode counting and unique molecular identifier (UMI) counting. Next, a digital gene expression matrix (gene counts versus cells) was generated. For quality control, we removed the cells with <200 genes or >5,000 genes detected, as well as cells where >15% of the transcripts were attributed to mitochondrial genes. The gene count matrix was converted to a Seurat object by the Seurat R package^37^. We normalized the filtered gene expression matrix by the *NormalizeData* function in Seurat R package, in which the number of UMIs of each gene was divided by the sum of the total UMIs per cell, multiplied by 10,000, and then transformed to log-scale.

### Dimension reduction and osteoblastic clusters identification

For data visualization and classification, we projected the normalized gene expression matrix on a two-dimensional panel. The 2,000 genes with the highest dispersion (variance/mean) were selected for principal component analysis (PCA). The first 18 principal components (number of PCs was chosen based on standard deviations of the principal components, corresponding to the plateau region of ‘‘elbow plot’’) were used for uniform manifold approximation and projection (UMAP)^36^ dimension reduction. After data visualization, we applied an unbiased graph-based clustering method^89^ for clustering analysis. To identify the differentially expressed genes (DEGs) in each cluster, we used the Wilcoxon Rank-Sum test to find the genes showing significantly higher levels of expression (false discovery rate (FDR) < 0.05) in a specific cluster compared to the other clusters.

### Pathway enrichment and trajectory inference analysis

To investigate the biological processes and signaling pathways associated with each subtype, we performed gene ontology (GO) and Kyoto encyclopedia of genes and genomes (KEGG) enrichment analysis for the genes identified as important cluster DEGs (avg_FC >1.2000) by using the *clusterProfiler* R package^90^. We then used the *Monocle 2 v2.8.0*^9^ R package to reconstruct the single-cell developmental trajectories in pseudo-time order. The principle of this analysis is to determine the pattern of the dynamic process experienced by the cell population and to order the cells along their developmental trajectory based on differences in the expression profiles of highly variable genes.

### Cross-species scRNA-seq data integration

One previous scRNA-seq dataset of mouse osteoblasts was acquired from GEO database under the accession numbers of GSE108891^31^. After acquiring the expression matrix of the osteoblasts in mice, we clustered them into three subtypes (M1, M2, and M3), following the analysis pipeline proposed by Tikhonova et al.^31^. Next, we integrated the scRNA-seq data of mouse and human osteoblasts by canonical correlation analysis (CCA) using Seurat R package^92^. The biological variance of transcriptional profiles across humans and mice was then evaluated based on the Spearman correlation of average gene expression between each of the datasets.

### Bone sectioning, immunostaining, and confocal imaging

Freshly dissected femur from female C57BL/6 mouse was fixed in 4% paraformaldehyde overnight followed by decalcification in 10% EDTA for 1 week. Samples were cut in 5-μm-thick longitudinally oriented sections. After deparaffinize and antigen retrieval, sections were blocked in PBS with 5% bovine serum albumin (BSA) for 1 hour and then stained overnight with goat-anti-Alpl (R&D: AF2910-SP, 10 μg/mL). Rabbit-anti-Nur77 (Proteintech:12235-1-AP, 1:2000) was used as secondary antibodies (from Invitrogen, 1:400). Slides were mounted with anti-fade prolong gold (Invitrogen) and images were acquired with a Zeiss LSM780 confocal microscope.

### Osteogenesis induction

Murine MSCs (1.0 × 10^4^ per well) were plated in 48-well plates and cultured in MesenCult™ basal expansion medium with 10% 10x Supplement (Stemcell) for 72 h. Next, the cells were rinsed with PBS and the medium was replaced with osteogenic differentiation medium (Stemcell). MSCs cultured in expansion medium were served as the negative control. Half of the medium was changed every 3 days, and cells were harvested at 0, 5, 7 days after induction.

### qRT-PCR analysis

Total RNA was extracted using an RNA extraction kit (Qiagen, Hilden, Germany). and cDNA was synthesized from 1 μg of total RNA by using the Revert Aid First Strand cDNA synthesis kit (Thermo). Then, the cDNA was amplified with iTaq™ Universal SYBR^®^ Green Supermix (BioRad, Hercules, CA). in an ABI PRISM^®^ 7900HT System (Applied Biosystems, Foster City, USA). The relative standard curve method (2^−ΔΔ^CT) was used to determine the relative gene expression and GAPDH was used as a housekeeping gene for internal normalization. The PCR primers used in this study were as follows:

GAPDH: forward, 5 ‘-CACCATGGAGAAGGCCGGGG-3’, reverse, 5’-GACG- GACACATTGGGGGTAG-3’;
ALP: forward, 5’-CCAACTCTTTTGTGCCAGAGA-3’, reverse, 5 ‘-GGCTACATTGGTGTTGAGCTTTT-3’;
NR4A1: forward, 5 ‘-AGGGTGTGTGTGCATATGGA-3’, reverse, 5 ‘-CCGCCATCTTTTCCTGTACG-3’;

