## Supplementary figures and images for "A systematic dissection of human primary osteoblasts *in vivo* at single-cell resolution"

### Figure_S1.pdf

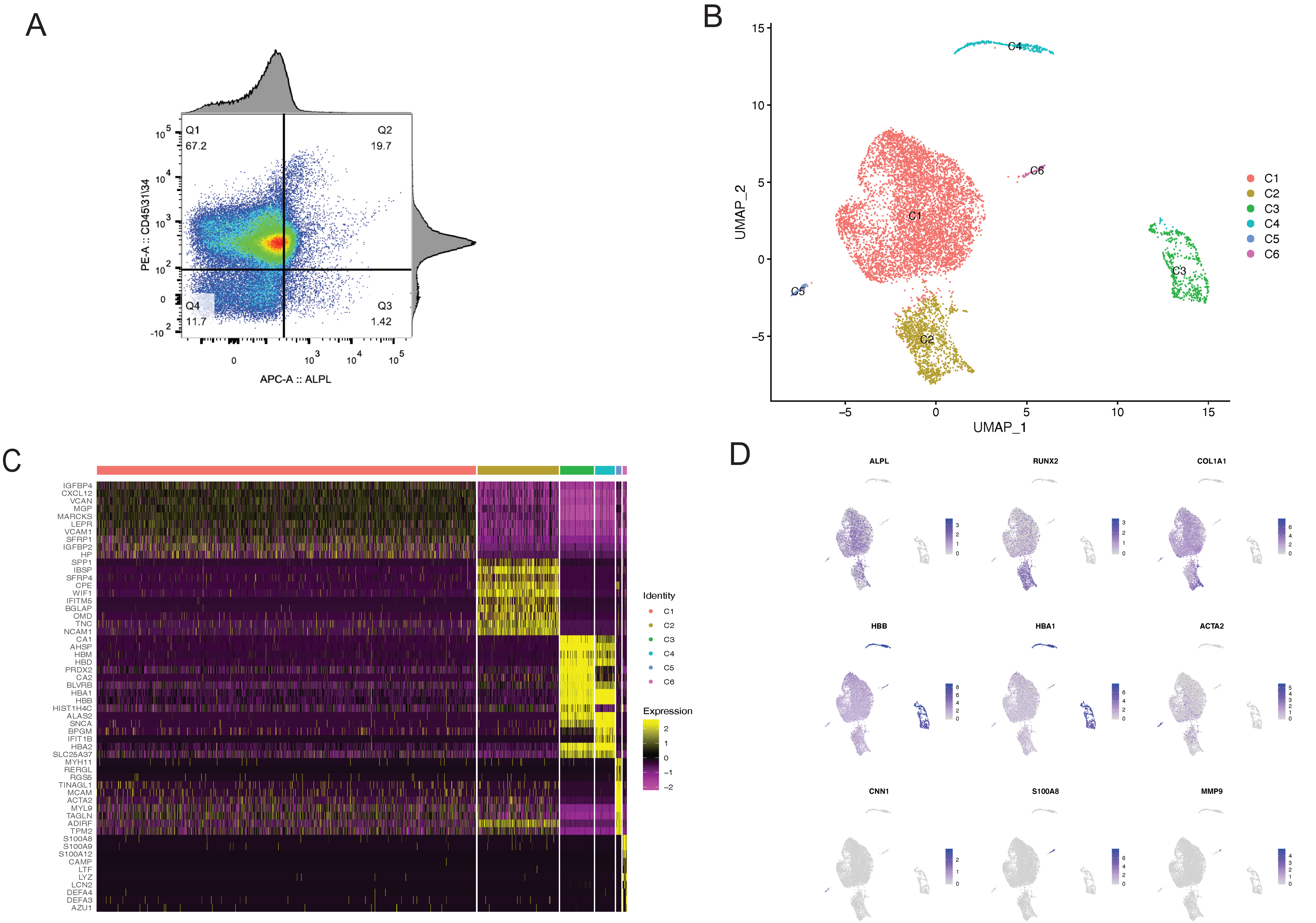
