## Supplementary material for "A systematic dissection of human primary osteoblasts *in vivo* at single-cell resolution": Supplemental information (1)-D.docx

**Supplemental Tables**

**Table S1.** DEGs in osteoblasts (clusters C1 and C2) compared with other contamination cells (clusters C3, C4, C5, and C6). The p_val and p_val_adj indicate the p-value and p-value adjusted by FDR. The avg_FC column in the table represents the average fold change of gene expression.

**Table S2.** DEGs with avg_FC > 1.200 in each cluster compared with other clusters.

**Table S3-S5.** Enriched GO terms for osteoblast subtypes O1, O2, and O3, respectively.

**Supplemental Figure Legends**

**Figure S1. Osteoblasts isolation and identification**

(A) Representative flow cytometry of osteoblasts isolation. The x-axis represents the expression level of ALPL and the y-axis is the expression level of CD45/31/34. Dots represent the cells. Cells in Q3 panel (downer-right) are isolated for downstream analysis.

(B) UMAP dimension reduction of isolated cells, colored by different clusters.

(C) Heatmap of gene expression profile of isolated cells, based on the relative gene expression level of top 10 most-significant markers for each cluster. X-axis represents different osteoblast clusters, while y-axis indicates the 10 most-significant markers in each cell cluster.

(D) Expression of known cell markers of each isolated subpopulations embedded on UMAP dimension reduction map. The first three markers (ALPL, RUNX2 and COL1A1) in the first row are the osteogenesis markers. HBB and HBA1 are markers of two nucleated erythrocyte clusters C2 and C3. The last four genes are markers for cluster C4 and C5, respectively.
